# OmicPioneer-sc: an integrated, interactive visualization environment for single-cell sequencing data

**DOI:** 10.1101/2020.10.31.363580

**Authors:** John N. Weinstein, Mary A. Rohrdanz, Mark Stucky, James Melott, Jun Ma, Vakul Mohanty, Ganiraju Manyam, Christopher Wakefield, Ken Chen, Nicholas E. Navin, Michael C. Ryan, Rehan Akbani, Bradley M. Broom

**Affiliations:** Department of Bioinformatics and Computational Biology, University of Texas MD Anderson Cancer Center, Houston, TX 77030, U.S.A; Quantitative Research Computing, University of Texas MD Anderson Cancer Center, Houston, TX 77030, U.S.A; In Silico Solutions, Fairfax, VA 22031, U.S.A; Department of Genetics, University of Texas MD Anderson Cancer Center, Houston, TX 77030, U.S.A

**Keywords:** Next-Generation Clustered Heat Map, NGCHM, clustered heat map, t-SNE, UMAP, pathway, single-cell sequencing, single-cell RNA-seq, visualization, bioinformatics, genomics, PathwayMapper

## Abstract

OmicPioneer-sc is an open-source data visualization/analysis package that integrates dimensionality-reduction plots (DRPs) such as t-SNE and UMAP with Next-Generation Clustered Heat Maps (NGCHMs) and Pathway Visualization Modules (PVMs) in a seamless, highly interactive exploratory environment. It includes fluent zooming and navigation, a statistical toolkit, dozens of link-outs to external public bioinformatic resources, high-resolution graphics that meet the requirements of all major journals, and the ability to store all metadata needed to reproduce the visualizations at a later time. A user-friendly, multi-panel graphical interface enables non-informaticians to interact with the system without programming, asking and answering questions that require navigation among the three types of modules or extension from them to the Gene Ontology or information on therapies. The visual integration can be useful for detective work to identify and annotate cell-types for color-coding of the DRPs, and multiple NGCHMs can be layered on top of each other (with toggling among them) as an aid to multi-omic analysis. The tools are available in containerized form with APIs to facilitate incorporation as a plug-in to other bioinformatic environments. The capabilities of OmicPioneer-sc are illustrated here through application to a single-cell RNA-seq airway dataset pertinent to the biology of both cancer and COVID-19.

[Supplemental material is available for this article.]

## Introduction

No single type of visualization suffices to capture all useful aspects of large omic datasets such as those generated by single-cell sequencing (or flow cytometry). The visual most frequently used over time has been the Clustered Heat Map (CHM), which we introduced in the early 1990s to capture patterns in large omic datasets (Weinstein et al. 1994; Weinstein et al. 1997). More recently, we have developed highly interactive “Next-Generation Clustered Heat Maps” (NGCHMs) (Broom et al. 2017; Ryan et al. 2019), which enable the user to navigate and zoom fluently from global to detailed views of the map, invoke a statistical toolbox, link to dozens of external public bioinformatic resources, produce high-resolution graphics, and save all metadata needed to reproduce the map even months or years later. The novel NGCHM technology is central to OmicPioneer-sc.

Particularly in single-cell studies, a second, widely used class of visuals is what we will call here generically a “Dimensionality Reduction Plot” (DRP). DRPs are 2D (or 3D) scatter plots in which each point represents a single cell whose position in the plot is determined by a vector of values representing some set of cell features, e.g., the cell’s scRNA-seq transcriptomic profile across genes. Classical linear DRP algorithms used in single-cell analyses have included Multidimensional Scaling (Urpa and Anders 2019) and Principal Components Analysis (Wold et al. 1987). More recently, non-linear algorithms like t-SNE (Maaten et al. 2008) and UMAP (McInnes et al. 2018), which more clearly delineate the differences among cell types, have been widely used. A third type of visualization is what we refer to generically as a Pathway Visualization Module (PVM), which most frequently takes the form of a functional cell pathway, cell network, or Gene Ontology.

OmicPioneer-sc integrates DRPs, NGCHMs, and PVMs in a seamless, highly interactive exploratory environment that captures the capabilities of all three of those versatile visualization/analysis paradigms. A user-friendly graphical interface enables non-informaticians to explore the visualizations, asking and answering questions that require navigation among them. Key features of OmicPioneer-sc include a high degree of user-friendly interactivity, immediate access to dozens of external tools and data resources, and inclusion of all 3 types of visualization, with the capability to add more in the future. To name one use-case, inclusion of PVMs can aid in the detective work required for identification and annotation of cell-types in the DRP and NGCHM. The ability to navigate interactively among cell, sample, feature, and pathway levels of information can also help address the complex molecular heterogeneity that often makes it challenging to isolate interesting cell phenotypes (Tirosh et al. 2016; Izar et al. 2020) or analyze cell functional status (Liang et al. 2020)

All-in-one analysis software, including Seurat (Butler et al. 2018) and SimpleSingleCell in Bioconductor (Lun et al. 2016), have been developed for processing and analysis of scRNA-seq data. However, they focus more on the data pre-processing and analysis steps (batch correction, differential expression analysis, clustering), and they require programming, typically in R. Unlike OmicPioneer-sc, however, they do not provide a navigable, dynamic environment for visualization of the data or for linking-out to external information resources. Likewise, commercial analysis packages for scRNA-seq data analysis (e.g., 10X Genomics Cell Loupe) have limited interactive features beyond clustering of cells and identification of the most highly expressed genes in each cluster. There remains a need in the field for highly interactive, user-friendly, web-based visualization-oriented software like OmicPioneer-sc for analysis and interpretation of scRNA-seq data and other types of data from single-cell studies.

## Results

Fig. 1 shows a conceptual schema for the OmicPioneer-sc system as a whole. The user can, for example, select a set of cells in the DRP, thereby automatically selecting them also in the NGCHM and in a salient PVM for enrichment calculations. Alternatively, the user can start with a particular PVM and project the relevant genes into the extensive resources of the NGCHM module or into the DRP. Note that DRPs submerge the identities of individual genes, and PVMs submerge the identities of individual cells, whereas NGCHMs capture both cell and gene information explicitly (in zoomable, navigable form). In the next section, we briefly describe each of the three module types and then show how they are integrated to form the overall system.

**Fig. 1.**
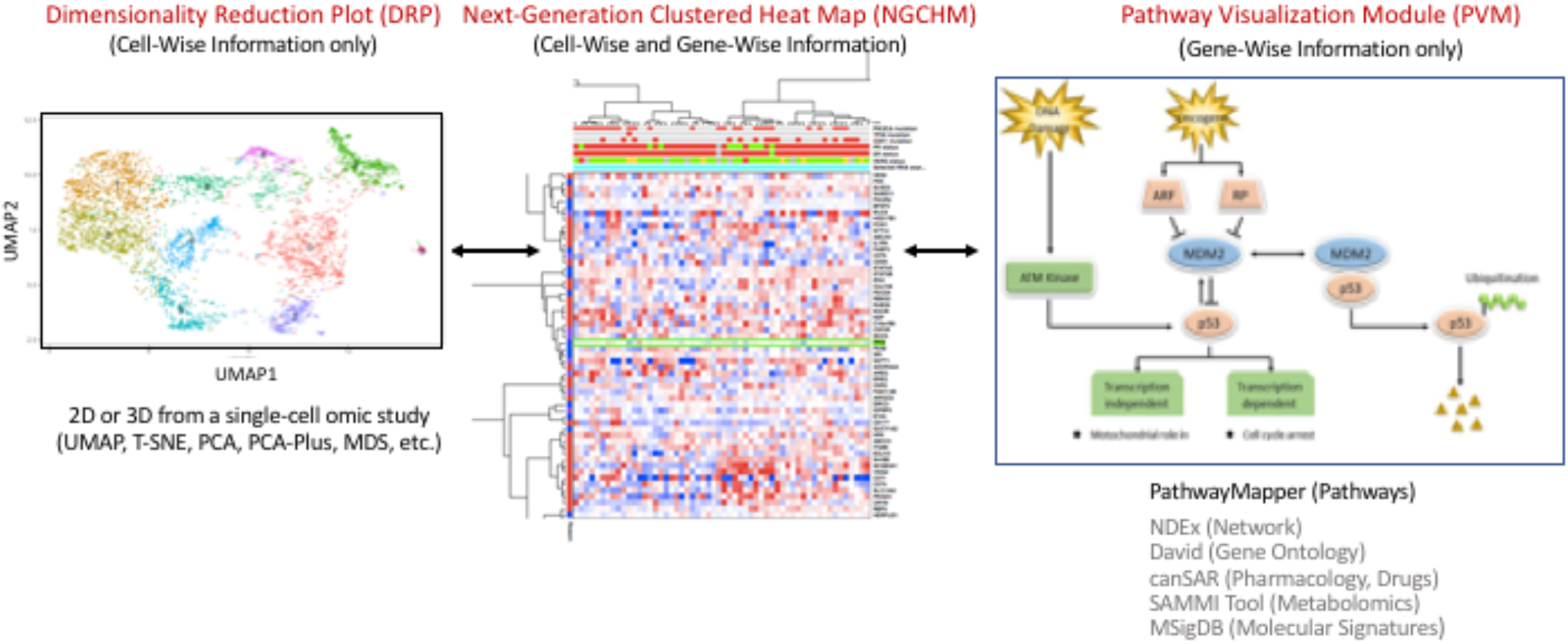
Conceptual schematic view of OmicPioneer-sc’s capabilities illustrated in terms of a 2D UMAP, an NGCHM, and a pathway diagram. DRPs submerge information on individual genes; PVMs (pathways, networks, ontologies) submerge information on individual cells; NGCHMs present both types of information in explicit form. Modules listed in grey text are planned or in development but not implemented in the current publicly available software.

For concreteness, the descriptions here will assume input data in terms of genes and single cells (e.g., data from scRNA-seq, snRNA-seq, or ATAC-seq at the gene level). For most purposes, however, one could substitute “sample” for “cell” or substitute another datum type such as “DNA methylation probe” or “ATAC-seq sequence” for “gene.” Link-outs obviously depend on there being some type of stereotyped unique identifier (e.g., HUGO gene names in the case of gene-level data). The same generic structure can be optimized for application to non-sequencing datasets such as those from flow cytometry or studies of bulk tissue or tumor.

We will use the term “TriptOme” (loosely derived from “triptych of omic data”) to denote a specific instance of the three visualizations consisting of DRP, NGCHM, and PVM. The OmicPioneer-sc suite of software tools centers on the NGCHM Viewer, which provides the overall user interface, displays the NGCHM, provides the link-outs, and closely integrates with the other visualization tools, including the PVM and DRP modules. The suite includes tools for creating TriptOmes, as well as servers for hosting databases and large compendia of the visualizations. A prebuilt version of the TriptOme described in this paper (Fig. 2) can be viewed and interactively explored online (without programming or entry of a database) at https://link.ngchm.net/FLNYf6Mxr (with an explanatory video at https://link.ngchm.net/zmwup-8UP).

**Fig. 2.**
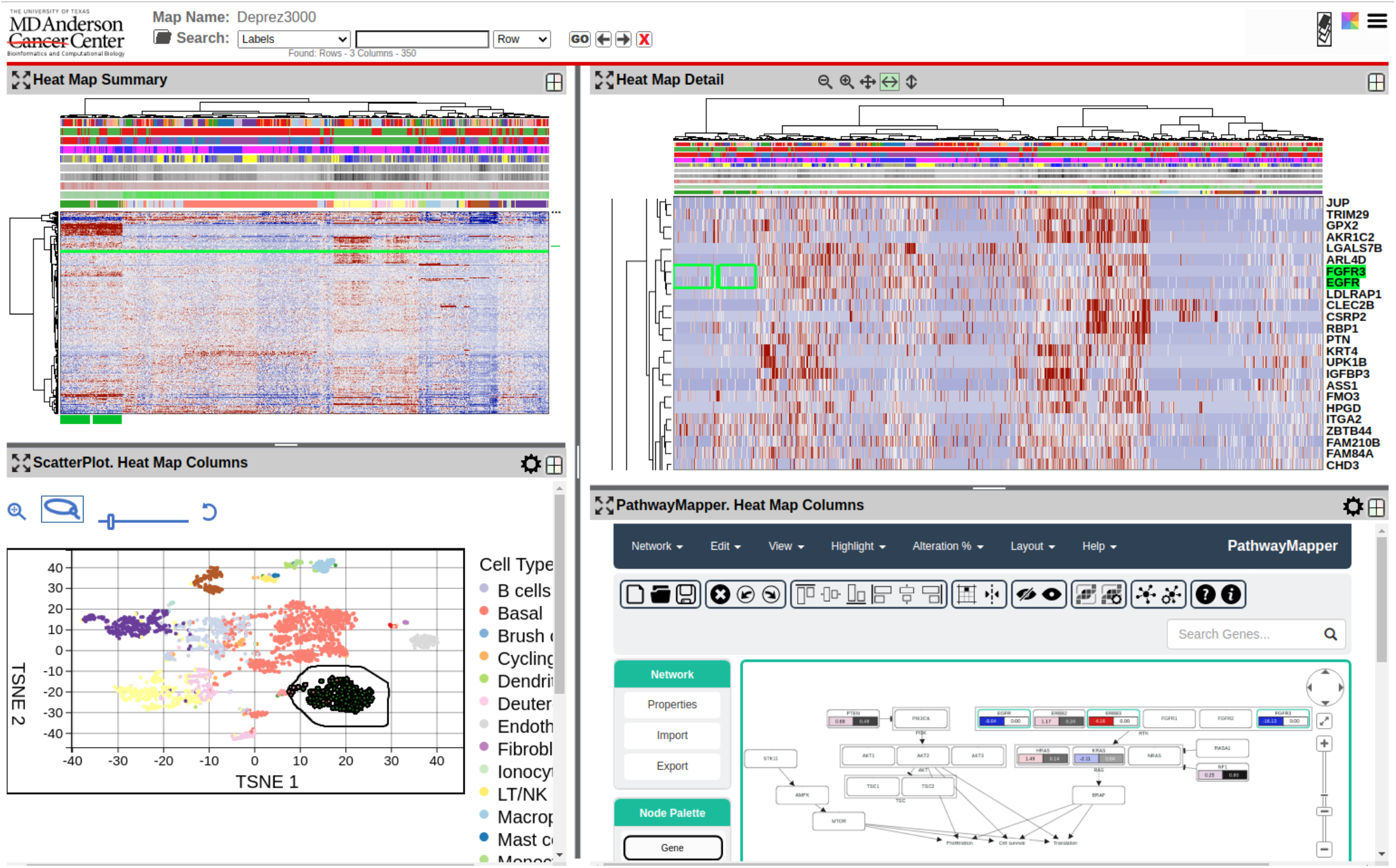
Screen shot of a four-panel OmicPioneer-sc visualization. **Upper left:** Global view of an interactive NGCHM of singlecell human lung RNA-seq data (Ward’s clustering algorithm, Euclidean metrics on both axes). There are 3,000 rows (genes) and 3,000 columns (cells) in this relatively small-scale example. **Upper right:** Zoomed view of one small portion of the NGCHM. **Lower left:** t-SNE plot of the data. Data points (cells) are colored according to the Cell Type covariate bar in the NGCHM (i.e., the bar closest to the heat map). Lasso and navigation tools are at the upper left. The lasso has been used to select the cluster of cells inside the black boundary, and those cells are then also marked (green rectangles) in the NGCHM global and zoomed views. **Bottom right:** A PathwayMapper panel showing RAS pathway genes, with color-coded numbers generated by a t-test comparing the selected cells with all others. The two most negative (blue) entries are highlighted in the Zoomed view (top right). Data are from (Deprez et al. 2020), which presents a detailed single-cell atlas of the human airway from nose to the 12^th^ division of the airway tree. Single-cell capture was performed using the 10X Genomics Chromium device (3’ V2), and data from the 35 samples for the atlas were pre-processed (Deprez et al. 2020) using fastMNN (Haghverdi et al. 2018) and Scanpy (Wolf et al. 2018). We used the pre-processed scRNA-seq data and metadata in OmicPioneer-sc to generate the images shown. The Supplemental Methods section includes the code used to generate the figure, along with an example NGCHM, and detailed explanations. The dataset provides a baseline for study of pathological and immunological changes in lung tissue such as those that can arise in carcinogenesis or viral infection (e.g., by SARS-CoV-2).

Text and figures that follow in this Results section explain many of the features that can be tried out in the pre-built version. For example, selecting a gene or genes (adjacent or disjoint) and then right-clicking (control-clicking on a Mac) brings up a menu with the dozens of link-out choices. More comprehensive information and instructions for R-scripted implementations are available in the form of user guides, introductory videos, and tutorial videos listed (with links) in Table 1 in the Implementation and Availability section. Additional details can be found in the Supplemental Methods.

**Table 1.** Links to online OmicPioneer-sc and NG-CHM resources.

| Description | URL |
| --- | --- |
| 1. The main NGCHM project page. | https://bioinformatics.mdanderson.org/public-software/ngchm/ |
| **Videos (YouTube):** |  |
| 2. Explanatory video of OmicPioneer-sc | https://link.ngchm.net/zmwup-8UP |
| 3. Select NG-CHM Demo Videos. | https://www.ngchm.net/demo-video |
| 4. A playlist of tutorial videos on constructing NGCHMs using R. | https://www.youtube.com/playlist?list=PLIBaINv-Qmd0v2qD4AMbGf85Xd-T_yzEh |
| 5. Video that steps through the tutorial script (included in row 4 playlist). | https://link.ngchm.net/UJxTTwMRC |
| **On-line Resources:** |  |
| 6. Our compendium of hundreds of NGCHMs on data from The Cancer Genome Atlas and PanCancer Atlas programs. | https://bioinformatics.mdanderson.org/TCGA/NGCHMPortal/ |
| 7. View the Triptome described in this paper (Fig. 2) interactively. | https://link.ngchm.net/FLNYf6Mxr |
| **Downloads:** |  |
| 8. Downloadable copy of the Triptome described in this paper. Open with the viewer (next row). | https://link.ngchm.net/YnV2kgbCv |
| 9. A downloadable version of the NG-CHM viewer. (By default, clicking the link may open the NG-CHM Viewer in the browser. Alternatively, you can save the link contents to disk and open it at a later time.) | https://www.ngchm.net/Downloads/ngChmApp.html |
| 10. A detailed knitr script for creating the TriptOme described in this paper. | https://www.ngchm.net/Downloads/how-to-create-single-cell-ngchm-in-r |
| **Source Code Repositories:** |  |
| 11. Core NGCHM module | https://github.com/MD-Anderson-Bioinformatics/NG-CHM |
| 12. Module for 2D DRP | https://github.com/MD-Anderson-Bioinformatics/ScatterPlotPlugin |
| 13. Customized PathwayMapper PVM | https://github.com/MD-Anderson-Bioinformatics/pathway-mapper |
| 14. NGCHM R package | https://github.com/MD-Anderson-Bioinformatics/NGCHM-R |

### Panel structure

Fig. 1 shows the conceptual framework of a TriptOme as a linear progression from DRP to NGCHM to PVM. However, to provide flexibility and optimize use of screen space, the OmicPioneer-sc viewer displays the modules in a set of panels that the user can select and configure as desired. Fig. 2 shows a four-panel arrangement consisting of a t-SNE plot, a global-view NGCHM, a detailed-view (i.e., zoomed) NGCHM, and a PathwayMapper panel. The relative dimensions of the various panels can be adjusted by the user, and the panels can be split vertically or horizontally to accommodate additional panels if desired. The icon at the upper right of each panel opens a dropdown menu that contains multiple options for the panel. For example, the menu could be used to open a fifth panel to show a UMAP for direct comparison with the t-SNE plot. At the upper left of each panel is a button that zooms the panel to occupy the full window for maximum visibility.

### Data pre-processing

We recognize that users will generally have their own preferred tools and methods for clustering and for data preprocessing functions such as filtering, preliminary dimensionality reduction, and batch effect correction, as well as imputation of values for dropouts (Andrews and Hemberg 2018; Peng et al. 2019). For example, Tran et al. (2020) have benchmarked 14 different algorithms for batch effect correction in single-cell studies: MNN Correct (Haghverdi et al. 2018), fastMNN (Haghverdi et al. 2018; Lun 2019), MultiCCA Seurat 2 (Butler et al. 2018), Seurat 3 (Stuart et al. 2019), MMD-ResNet (Shaham et al. 2017), Harmony (Korsunsky et al. 2019), Scanorama (Hie, et al. 2019), BBKNN (Polański et al. 2020), scGen (Lotfollahi et al. 2018), ComBat (Johnson et al. 2007), LIGER (Welch 2018), limma (Smyth and Speed 2003), scMerge (Lin et al. 2019), and ZINB-WaVE (Risso et al. 2018). Li et al. (2020) used an unsupervised deep-learning algorithm for simultaneous clustering and batch effect correction. Novel approaches to pre-processing and clustering, with the almost bottomless diversity of important nuances and special cases that those processes entail (see, e.g. Kiselev et al. 2019), are not our aim here; OmicPioneer-sc’s novelty is focused downstream on interactive visualization and interpretation of patterns in the data. Cell-type annotations generated using any automated or semi-automated method, including R packages such as SingleR (Aran et al. 2019), CellAssign (Zhang et al. 2019), and DESC (Li et al. 2020), can be attached as covariate bars and used for selection of colors and/or point types in the DRP module. Integration of PVMs with the other OmicPioneer-sc modules can aid in the detective work required for the functional annotation process.

### The Dimensionality Reduction Plot (DRP) module

The DRP module’s visualization code is agnostic in that the input data and visualization machinery are the same for any 2D (or 3D) scatter plot, regardless of the algorithm that generated it (e.g., Principal Component Analysis, Multidimensional Scaling, t-SNE, UMAP). Hence, the user can run any favorite package to generate the cell coordinates for the DRP. Once the set of coordinates plus cell-type identifications have been provided, the user can access the TriptOme’s customized DRP viewer. The viewer is full-featured, including:

- Fluent zooming and navigation
- Flexible coloring of cell subsets on the basis of genes or signatures
- Capture of individual cells or multiple individual cells by clicking
- Selection of groups of cells (contiguous or not) using a free-form lasso
- Flexible control of cell symbols in the figure, including adjustment of cell symbol size for optimal viewing
- Zooming to full-window size for enhanced visibility

As a default, cell symbols are drawn in random order to prevent visual bias toward the last colors applied if there are overlapping data symbols. However, the user can choose interactively to draw and color cell types in order from largest to smallest cell group to avoid masking the latter if the symbols do overlap. As noted, the DRP coordinates can be calculated using any state-of-the-art algorithm, including an initial dimensionality reduction using principal components analysis, independent components analysis, or another such method, as reviewed in Sun et al. (2019). The resulting dimensionality summary data (RDSD) can be attached to the NGCHM as an auxiliary data set for use in additional analyses. The most significantly novel aspect of the DRP viewer is its customization for interactive, automated syncing with the NGCHM, which will be described in the next section.

### The Next-Generation Clustered Heat Map (NGCHM) module

The NGCHM module’s capabilities are detailed in our previous publications (Broom et al. 2017; Ryan et al. 2019), in a series of YouTube videos, and in a User Guide (see Table 1 for details). Briefly: In contrast to the early, static CHMs (Weinstein et al. 1994; Weinstein et al. 1997), NGCHMs use a dynamic tiling strategy to enable fast, fluent, extreme zooming from a global view of the map to detailed views in which the gene and/or cell labels can be displayed visibly. If the labels are HUGO gene names, a context-sensitive menu with links to more than two dozen external resources appears (as illustrated in our compendium of hundreds of NGCHMs on data from The Cancer Genome Atlas and PanCancer Atlas programs (see Table 1, row 6)). Analogously, if the data are from reverse-phase proteomic arrays (RPPA), the labels (RRIDs) and their link-outs correspond to the antibodies used for detection of proteins on the arrays. Our NGCHMs on TCGA microRNA datasets link to miRBase, among other resources. Data at the subgene level from technologies such as scATAC-seq and single-cell DNA methylation studies can currently link to generic external resources and will, in the near future, link to pertinent specialized resources. Other features of NGCHMs include interactive re-coloring on the fly, multiple methods for selecting sets of genes or cells in the map (including from the dendrograms), flexible search functions, and many controls for customizing the dimensions and “look” of the figure (e.g., the font type and size). A statistical toolkit and an ideogram viewer linked to the UCSC Genome Browser (Lee et al. 2020) can be invoked. Output options include the original data, high-resolution pdfs, and metadata sufficient to reproduce and/or share the map at a later date. Overall the interactive NGCHM package is considerably fuller-featured than any other CHM viewer of which we are aware. We have implemented plug-in status or links with a number of capable bioinformatic environments, including cBioPortal (Cerami et al. 2012), GenePattern (Reich et al. 2017), Galaxy (Afgan et al. 2018; Broom et al. 2017), Trinity (Grabherr et al. 2011), and the Metabolomics Workbench (Sud et al. 2016). That has often been done through APIs generated in collaboration with the pertinent partnering development team.

In the single-cell context, RDSD data from the DRP computation can be input to the hierarchical clustering algorithm used to generate the dendrogram for the cell axis of the NGCHM. Alternatively, cluster assignments calculated using a graph-based (instead of hierarchical) clustering algorithm can be attached as covariate bars to the NGCHM and no dendrogram displayed. Our experience to date has shown a considerable degree of concordance among different cluster assignment methods. Genes that differ in expression significantly among clusters can be indicated using a covariate bar attached to the gene (row) axis of the NGCHM. The NGCHM’s search function enables such genes to be located rapidly and used for coloring a DRP. The module enables the user to determine at a glance whether all of the genes concerned are highly correlated or belong to two or more disjoint groups.

### The Pathway Visualization Module

The PVM enables the user to project data from the DRP through the NGCHM to pathways (or the reverse), for example to identify pathways enriched with genes characteristic of a given cell subset. Many different pathway paradigms can be accommodated, but the capable one implemented currently is a customized version (scPathwayMapper) of the popular, versatile PathwayMapper (Bahceci et al. 2017). A pathway diagram can be created interactively or loaded into the module from any of many sources, including the cBioPortal’s database of TCGA-related pathways and personal (file system) archives. The user can click on one or more genes in the pathway diagram (or click on all of them) to highlight them in the NGCHM and apply any gene-centric capability of OmicPioneer-sc (e.g., coloring of a DRP). The user can also color genes in the scPathwayMapper diagram using data from the NGCHM. For instance, two groups of cells can be selected using the DRP module’s lasso tool and colored in the pathway diagram using the results of a t-test between the two groups. Gene Ontology enrichment calculations (e.g., GSEA; Tamayo et al. 2005) and additional PVM types are planned as well. OmicPioneer-sc is under continuing, active development.

### Implementation and availability

OmicPioneer-sc is publicly available as an open-source project. Table 1 includes links to online resources, including, as noted earlier, a pre-built version of the TriptOme described in this paper (Fig. 2) that can be viewed and interactively explored online (Table 1 row 7), along with an explanatory video (Table 1 row 2). Alternatively, a stand the data files can be downloaded (Table 1 row 8) and viewed using the downloadable version of the NGCHM viewer (Table 1 row 9). The main NGCHM project page (Table 1 row 1) presents details of the NGCHM module. Our GitHub site includes the following open-source projects:

- the NGCHM module (Table 1 row 11);
- a customized 2D DRP module (Table 1 row 12);
- a customized PathwayMapper PVM (Table 1 row 13).

Currently, TriptOmes can be created using the NGCHM R package (Table 1, row 14) in conjunction with common R data structures or packages used in single cell analysis, including singleCellExperiment (Lun et al. 2020), Seurat (Stuart et al. 2019), and singleCellToolkit (Jenkins 2019). The NGCHM R package allows basic NGCHMs to be created very easily, while enabling the user to augment or replace default values or algorithms with custom choices, including specialized R algorithms of arbitrary sophistication. The package also includes features that simplify the creation of advanced features or specialized variants of complete TriptOmes. Included are utility functions that can attach DRP coordinates from common PCA, UMAP, t-SNE, and singleCellExperiment data structures to an NGCHM while it is being created. For example, the following R statement attaches pre-computed UMAP coordinates from a singleCellExperiment object to the columns of an NGCHM during its construction:

~~~
hm <-chmAddReducedDimension (hm, “column”, sce, “UMAP”);
~~~

In addition to its built-in methods for generating the NGCHM’s dendrograms, the NGCHM R package can use any R dendrogram or other ordering algorithm for either axis. For OmicPioneer-sc, we note that if RDSD data are used for generating the DRP plot(s), the same RDSD data can be used as input to the hierarchical clustering algorithm.

A detailed knitr (Xie 2014) script (unpublished) for creating the TriptOme described in this paper is available at our website (Table 1, row 10), as are the data required by that script. The code and text explanations are included in the Supplemental Material. Our YouTube channel includes a playlist of tutorial videos on constructing NGCHMs using R (<u>Table 1, row 4</u>), including one that steps through the above tutorial script (Table 1, row 5).

The YouTube tutorials use a customized RStudio (RStudio 2020) container (see our Downloads page for details). Three varieties of containers are available: one that includes just the NGCHM R package, one that also includes the Bioconductor core software (Jenkins et al., 2019), and one that further includes many of the common single-cell R packages cited above (including singleCellExperiment, Seurat, and singleCellToolkit) that can be used to program DRPs. Our GUI-driven NGCHM builders (web-based, Galaxy, and GenePattern) currently do not include scripts for generating the DRP coordinates; that capability for our web-based builder is in development.

### Performance evaluation

We have tested OmicPioneer-sc on datasets of up to 65,000 by 11,000 elements. That size matrix is large but still not sufficient to handle all of the cells in some current single-cell studies, which may involve reads from hundreds of thousands of cells per sample. The fundamental limiting factor is not the visualization machinery, but rather the clustering, particularly for conventional hierarchical clustering algorithms; the time required to create maps grows quickly (generally O(n^2^) with respect to the larger map dimension). The data from experiments with hundreds of thousands of cells can be randomly down-sampled or some other preliminary dimensionality reduction can be performed. For large maps, strategies to reduce clustering time (such as using RDSD data) and R’s fastcluster package (Müllner et al. 2013) have to be considered. Hence, we are exploring higher-performance clustering solutions, including parallel computing, GPU computing, graph clustering algorithms, and approximate computation methods. On current desktop workstations, creating maps with up to 5,000 rows or columns interactively is reasonable, and larger datasets can be clustered in batch mode for later interactive visualization. Very large datasets may take longer to load initially, but a key point is that online response thereafter (e.g., in zooming, navigating, and re-coloring) is very rapid, generally independent of size. The reason is that OmicPioneer-sc viewer’s tiling system requires detailed data for only the zoomed region being displayed plus its surround but requires only summarized (i.e., low-resolution) data for the rest of the global map.

### Comparison with other software

DRPs are ubiquitous in the single-cell literature, and, as noted above, there are many scripts and tools available for generating and viewing them. Although some interact with small, static CHMs, we know of no comparable DRP tool that is tightly integrated with an omic-scale, dynamic CHM. Since CHMs are also ubiquitous in the biological literature, it is not surprising that many scripts and tools have been developed for creating them. An extensive search of such tools indicated that most produce heat maps that are static, modestly interactive, and/or not scalable. The nearest in capabilities to NGCHMs that we have found are those created by ClusterGrammer (Fernandez et al. 2017), Java Treeview 3 (Keil et al. 2019), Morpheus (https://software.broadinstitute.org/morpheus), and MicroScope (Khomtchouk et al. 2016). None of them, however, have the NGCHM’s diverse array of important capabilities. In a survey of 60 software packages that provide clustering in scRNA analysis, for example, we found 20 that produce heat maps, only one with very extensive interactivity, and none that approach NGCHMs in exploratory features. Key capabilities that distinguish the NGCHM module from alternatives include choice of symmetric and asymmetric zooming, link-outs to dozens of external bioinformatics resources, incorporation of statistical tools, links to genomewise data visualizations like our ideogram viewer or the UCSC Browser, scalability to tens of thousands of elements per axis for the interactive visualization, ability to produce publication-quality graphics, semi-automated building of compendia, fluent controls for covariate bars, re-coloring on the fly, reproducibility, and shareability of interactive maps (e.g., by email or on the cloud) (see Table 1, row 3). None of the others have automated information flow among DRP, CHM, and PVM components. OmicPioneer-sc is an ongoing Agile development program that solicits suggestions and critique via its social media and github presence.

## Discussion

Visualization of large data sets has played a variety of important roles in the development and application of omic science (see, e.g., Weinstein et al., 1997; Nattestad et al. 2020). The OmicPioneer-sc suite provides a highly interactive, exploratory environment that combines and links features of three centrally important visualization paradigms for single-cell sequencing studies: Dimensionality Reduction Plots (e.g., PCA, t-SNE, UMAP), our interactive Next-Generation Clustered Heat Maps (NGCHMs), and Pathway Visualization Maps (e.g. PathwayMapper). Fluent online navigation within and among the three can facilitate the detective work required for functional annotation of the cells and consequent coloring of the graphics for single-cell studies. It also provides an extensive set of interactive tools for downstream visualization, analysis and interpretation of the data, including the ability to zoom in and magnify particular features corresponding to rare cell types that might otherwise be missed. OmicPioneer-sc provides a visualization/analysis toolset with which to pursue several of the “grand challenges” to single-cell studies set forth in Lähnemann et al. (2020). The present configuration is tuned to the idiosyncrasies of single-cell sequencing, but it can also be applied to technologies like spatially-dependent multi-parameter single-cell phenotyping and flow cytometry, as well as studies on bulk tissues. The tools are available in containerized form, with APIs to facilitate sharing or incorporation as a plug-in to other bioinformatics environments.

## Supplemental Methods

See Supplemental_Methods.pdf for the code and text explanations necessary to create single-cell NGCHMs. Also see the extensive set of introductory and instructional videos cited in the Implementation and Availability Section.

## Acknowledgements

The Michael & Susan Dell Foundation; The Mary K. Chapman Trust; NCI P30 CA016672 (CCSG Bioinformatics Shared Resource); NCI U24 CA199461 (ITCR); NCI U24 CA210949 (Batch Effects Genome Data Analysis Center); NCI U24 CA210950 (RPPA Genome Data Analysis Center); U01 CA235510 (NIH Metabolomics Common Fund, Quality control of metabolomic data). We thank C. Ron Bouchard and D. Blackburn for assistance with manuscript and references.

## Author Contributions (optional)

(to be considered later)

